# Shifting transmission risk for malaria in Africa with climate change: a framework for planning and intervention

**DOI:** 10.1101/797050

**Authors:** Sadie J. Ryan, Catherine A. Lippi, Fernanda Zermoglio

## Abstract

**Background:** Malaria continues to be a disease of massive burden in Africa, and the public health resources targeted at surveillance, prevention, control, and intervention comprise large outlays of expense. Malaria transmission is largely constrained by the suitability of the climate for *Anopheles* mosquitoes and *Plasmodium* parasite development. Thus, as climate changes, we will see shifts in geographic locations suitable for transmission, and differing lengths of seasons of suitability, which will require changes in the types and amounts of resources.

**Methods:** We mapped the shifting geographic risk of malaria transmission, in context of changing seasonality (i.e. endemic to epidemic, and vice-versa), and the number of people affected. We applied a temperature-dependent model of malaria transmission suitability to continental gridded climate data for multiple future climate model projections. We aligned the resulting outcomes with programmatic needs to provide summaries at national and regional scales for the African continent. Model outcomes were combined with population projections to estimate the population at risk at three points in the future, 2030, 2050, and 2080, under two scenarios of greenhouse gas emissions (RCP4.5 and RCP8.5).

**Results:** Geographic shifts in endemic and seasonal suitability for malaria transmission were observed across all future scenarios of climate change. The worst-case regional scenario (RCP8.5) of climate change places an additional 75.9 million people at risk from endemic (10-12 months) exposure to malaria transmission in Eastern and Southern Africa by the year 2080, with the greatest population at risk in Eastern Africa. Despite a predominance of reduction in season length, a net gain of 51.3 million additional people will be put at some level of risk in Western Africa by midcentury.

**Conclusions:** This study provides an updated view of potential malaria geographic shifts in Africa under climate change for the more recent climate model projections (AR5), and a tool for aligning findings with programmatic needs at key scales for decision makers. In describing shifting seasonality, we can capture transitions between endemic and epidemic risk areas, to facilitate the planning for interventions aimed at year-round risk versus anticipatory surveillance and rapid response to potential outbreak locations.

## Background

Malaria causes an estimated 435,000 deaths per year, with the majority of cases occurring in Sub-Saharan Africa, affecting children under 5 disproportionately [1]. Recent advances in reducing case burdens in sub-Saharan Africa through bed net distribution, household level spraying, and rapid clinical diagnostic and treatment responses appeared to slow down in 2017 and 2018, leaving reduction, and eradication goals unmet, and an estimated 219 million cases in 2018 [1]. The WHO reported that for 10 high burden African countries, there was an increase of 3.5 million cases in 2017 over the prior year. This stall in reduction was largely attributed to a stall in investments in global responses to malaria. The U.S. remained the single largest international donor in 2017, contributing $1.2 billion (39% of the overall investment); it is projected that roughly $6.6 billion annually by 2020 will be needed for the global malaria strategy, underscoring the importance of knowing how much and where to invest.

Geospatial modeling approaches provide a flexible framework in which to explore possible future scenarios of malaria risk as a function of changing climate [2]. Mordecai et al. introduced a mechanistic nonlinear physiological temperature-driven malaria transmission suitability model in 2013, via incorporating temperature dependent traits of boththe mosquito and parasite, based on laboratory data [3]. This demonstrated that transmissibility of malaria is constrained between 17-34C, which will therefore limit the spatial distribution of malaria on the landscape. In addition, this model updated the optimum temperature for malaria transmission from 31C to 25C, and the model was well validated using 40 years of field observation data matched to specific location month and temperature [3]. Temperature has also been shown to be an important predictor of incidence in many locations [4], and the potential effects of climate-induced temperature shifts as an impact on intervention and vector control efforts have been noted [5]. In previous work, we found that the top quantile of predicted transmission suitability from the Mordecai et al. model, that is, the top 25% of the transmission or *R*_*0*_ curve, best captured spatial and seasonal risk for Africa, from independent models of malaria risk prediction, based on statistical models of spatial case data from the Mapping Malaria Risk in Africa (MARA) and Malaria Atlas Project (MAP) projects [2,6–8].

Climate change threatens to the alter the nature of future malaria exposure across Sub-Saharan Africa [2,6,7]. Many countries with a high burden of malaria now have weak surveillance systems and are not well positioned to assess disease distribution and trends, making it difficult to optimize responses and respond to outbreaks [9]. To date, knowledge on how climate driven changes in malaria risk will manifest at regional and national scales is limited, though such knowledge is critical to designing responses. Changes in both the areas and populations exposed to malaria risk will necessitate adaptive responses to address them. To inform these responses, we explored six scenarios of changing suitability, aligned to potential management strategies to address the changing risks. We provide an updated view of climate-driven malaria shifts in Africa from the 2015 mapping paper by Ryan et al [2], using the newer IPCC AR5 climate change scenario framework, explicitly defining season length to align with policy language, and including a sub-continental approach, aligning changes to regional scale planning.

The goals of this study were to (1) identify new areas that will emerge as suitable for malaria transmission under different scenarios of change; (2) identify areas that may experience reductions in transmission suitability season length; and (3) provide an estimate of the human population at risk under each scenario. These are presented in the language of malaria seasonality risk, to align with surveillance and intervention targeting goals, and summarized as regional scale outcomes, broadly aligned with USAID’s planning scales, as the parent aid organization of much of the US investment in the global malaria strategy.

## Methods

### Malaria Transmission

The model for temperature-dependent malaria transmission presented in Mordecai et al. (2013) used this expression for *R*_*0*_, the basic reproductive rate of the disease, in order to account for the fitting of these rates to laboratory measurements:

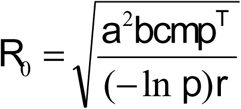

The temperature-dependent parameters are the mosquito biting rate (a), vector competence (b*c), mosquito density (m), the mosquito survival rate (p), and the parasite’s extrinsic incubation period (T), all of which are measurable empirical parameters.

The model incorporated temperature response curves fit for the mosquito species *Anopheles gambiae* and the malaria pathogen *Plasmodium falciparum*, with additional information used for related *Anopheles* and *Plasmodium* species. Transmission, *R*_*0*_ was scaled from 0–1, to describe relative transmission across the range of temperature. In Ryan et al [2] this curve was described this in quantiles, where the top quantile (upper 25 percent) of the curve was selected to represent the range of temperatures in which transmission suitability is expected. This conservative measure of the overall temperature curve was used as it corresponds to existing maps of ongoing transmission under current temperatures [2].

### Climate Data

Current temperature data is represented by globally gridded 5 arc-minute WorldClim (version 1) monthly mean temperature data [10]. This represents a long term average, or baseline, which has been used to project future climate scenarios, and therefore serves as our baseline.

General Circulation Models (GCMs) are the primary source of information about potential future climate. GCMs comprise simplified but systematically rigorous mathematical descriptions of physical and chemical processes governing climate, including the role of the atmosphere, land, oceans, and biological processes. They allow for modeling the expected climate response to increasing greenhouse gas concentrations. The direct application of GCM output to adaptation decision making, however, has been relatively limited due to GCMs’ coarse spatial resolution (100 to 500 km^2^). For strategic planning in malaria prevention and control, information is required on a much more local scale than GCMs can provide. Here, a statistically downscaled multi-model ensemble product is used for this analysis, compiled at a resolution of 5 arc-minutes (∼10 km^2^) from 6 downscaled GCMs. The climate projection data used in this study consisted of the median value for the multimodel ensemble representing future climate, compiled from the Coupled Model Intercomparison Project (CMIP5) archive, downscaled using a Change Factor (CF) approach and sourced from Navarro-Racines, Tarapues-Montenegro, and Ramírez-Villegas [11]. This ensemble approach allows exploration of the range of uncertainty across climate projections under two greenhouse gas emissions scenarios, or Representative Concentration Pathways (RCPs) – RCP 4.5 and RCP 8.5 – for three future time periods: the 2030s, 2050s, and 2080s.

### Aridity Masking

*Anopheles* mosquitoes (i.e., malaria-transmitting mosquitoes) require an appropriate level of moisture in their environment to provide breeding habitat with which to complete their lifecycle. Humidity or moisture is thus another component in the climate–transmission relationship. While several models use rainfall as a predictor for malaria occurrence, it is complicated to generalize how precipitation measures, such as monthly rainfall totals, cumulative rainfall, or relative humidity, actually manifest as breeding habitat for mosquitos at large scales [12–15]. Precipitation may not be a good indicator of standing water, and in a world of increasingly extreme precipitation events, the difference between a month’s rainfall occurring in a single day versus gradual accumulation over that month becomes more relevant. Mosquito habitat can wash away, “flushing” away eggs and disrupting the lifecycle, meaning that more rain does not necessarily translate into more habitat [16]. In addition, much of the world is subject to agricultural irrigation, redirecting precipitation in nonlinear ways at local level, or even creating piped water environments in the absence of precipitation. To generalize habitat suitability for mosquito breeding, a remotely sensed proxy is used: the normalized difference vegetation index (NDVI), which measures the photosynthetic activity of growing plant matter, on a 0-1 scale. The NDVI is thus a useful descriptor of the type of habitat conducive to *Anopheles* breeding. The threshold of “too dry” is based on prior work conducted by Suzuki et al. [17] to exclude locations where the NDVI drops below a critical minimum level for two months of the year, thereby cutting off breeding and the transmission cycle [17]. We followed a modified version of the methods of Ryan et al. [2] to limit projected models to those geographic areas capable of supporting mosquito survival. Monthly NDVI values were derived from post-processed MODIS data, available from FEWS-Net (Famine Early Warning System Network) [18]and month-to-month thresholding was calculated [17]. That is, if the NDVI value for two consecutive months fall below 0.125, it is assumed that an aridity boundary is crossed, indicating that that area (pixel) is considered too arid for malaria transmission to occur. We chose the 2016-2017 period of NDVI as an average climate year for the current decade. As NDVI cannot be projected into future scenarios, we use this as an average current aridity mask, which is a conservative approach.

### Population Data

We downloaded global gridded population products, the Gridded Population of the World (GPW), at a 30 arc-second (∼1 km^2^) resolution. Population data for Africa used as input for calculating population at risk (PAR) under the various transmission scenarios were derived from the Gridded Population of the World, Version 4 (GPWv4) [19], with baseline estimates derived from 2015 GPW data, while projected future populations were extracted from the 2020 layers.

### Geospatial projections of transmission

The gridded temperature data (current and future climate scenarios, month-wise) were constrained to the temperature range of the optimal quantile of transmission, and the resulting number of months of transmission suitability in each pixel recorded for all of Africa. The aridity mask was applied, and pixels falling in masked areas were given no value.

Seasons of transmission were defined based on the numbers of months of suitability, and criteria established by MARA were followed in defining malaria transmission suitability, with very slight additional granularity to better illustrate the impact of changing climate (Table 1).

**Table 1.** Definitions of malaria transmission suitability used in summarizing areas and population at risk.

| Malaria Suitability | Definition |
| --- | --- |
| Endemic | Malaria transmission suitability for 10-12 months of the year |
| Seasonal | Malaria transmission suitability for 7-9 months of the year |
| Moderate | Malaria transmission suitability for 4-6 months of the year |
| Marginal | Malaria transmission suitability for 1-3 months of the year |

In order to estimate the population at risk (PAR) for each geospatial research question, the suitability data were aggregated by a factor of 10 and aligned to the climate data, such that all analyses were conducted at 5 arc-minute resolution (approximately 10 km^2^ at the equator). Population data for each scenario were summarized by region, shown in Figure 1. We defined five regions of Africa; these align with the policy scale, but not definition of countries for USAID’s four African regions. We chose to delineate Eastern Africa and Central Africa to align with physical geography – while USAID defines Eastern Africa to include the Democratic Republic of Congo and Congo, and Central African Republic, Cameroon, Gabon and Equatorial Guinea are all included in the USAID West African Region, we chose to define a Central Africa region, comprising these countries (Figure 1). We present results of our analyses for four of our regions, excluding Northern Africa from this study.

**Fig. 1.**
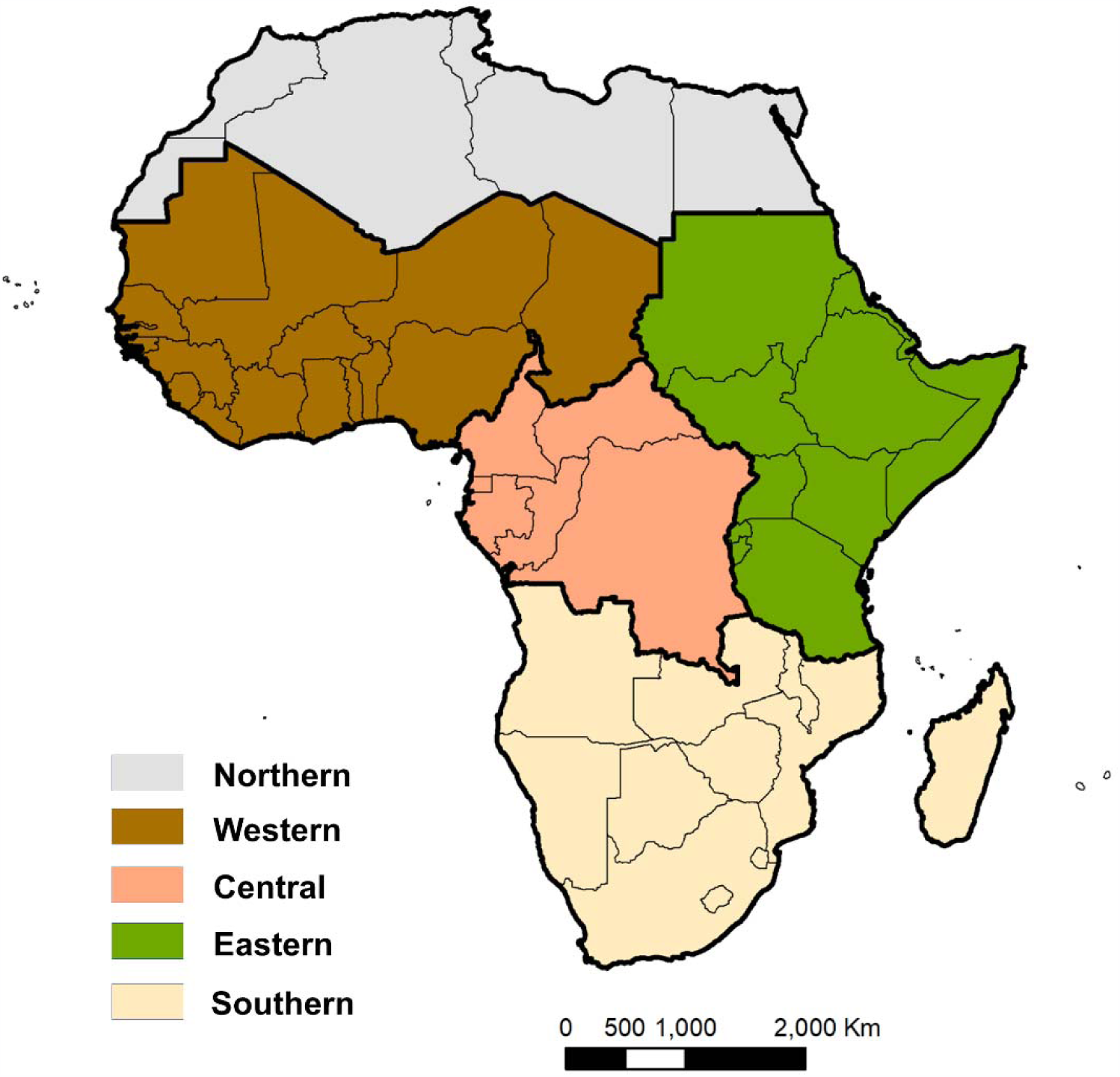
Map of the five regional definitions of Africa used in this study. Note that the Northern Africa region was excluded from analyses in this study.

All calculations and analyses were conducted in R [R version 3.3.3 2017-03-06 “Another Canoe”] using the “raster,” “rgdal,” “sp,” and “maptools” packages, and mapped output was produced in ArcGIS [Version 10.5.1].

## Results

### Regional impacts of climate change scenarios

Increases in temperature by region, from baseline, for the future climate scenarios, are synthesized in Table 2. Higher future temperatures are projected under all models and time periods evaluated for the continent.

**Table 2.** Average annual temperature increases (°C) from baseline (1960–1990) by region, RCP, and time period.

| Region | 2030s |  | 2050s |  | 2080s |  |
| --- | --- | --- | --- | --- | --- | --- |
|  | RCP 4.5 | RCP 8.5 | RCP 4.5 | RCP 8.5 | RCP 4.5 | RCP 8.5 |
| West Africa | 1.32 | 1.57 | 2.29 | 2.32 | 2.84 | 4.38 |
| East Africa | 1.32 | 1.63 | 1.90 | 2.32 | 2.96 | 4.38 |
| Central Africa | 1.10 | 1.42 | 1.63 | 2.07 | 2.69 | 4.04 |
| Southern Africa | 0.94 | 1.28 | 1.33 | 2.01 | 2.51 | 4.08 |

### Current and Future Suitability Risk

Under baseline conditions, we see the current distribution of endemic (10-12 months) transmission suitability for malaria is concentrated in the Central African region, with additional areas along the southern coast of Western Africa, and along the eastern coast of Eastern Africa, and in the north of Madagascar (Figure 2). Seasonal transmission (7-9 months of the year) suitability is predicted along a band through Western and Eastern Africa, south of the areas too arid for mosquito life cycles, and in parts of Southern Africa, particularly through Mozambique.

**Fig. 2.**
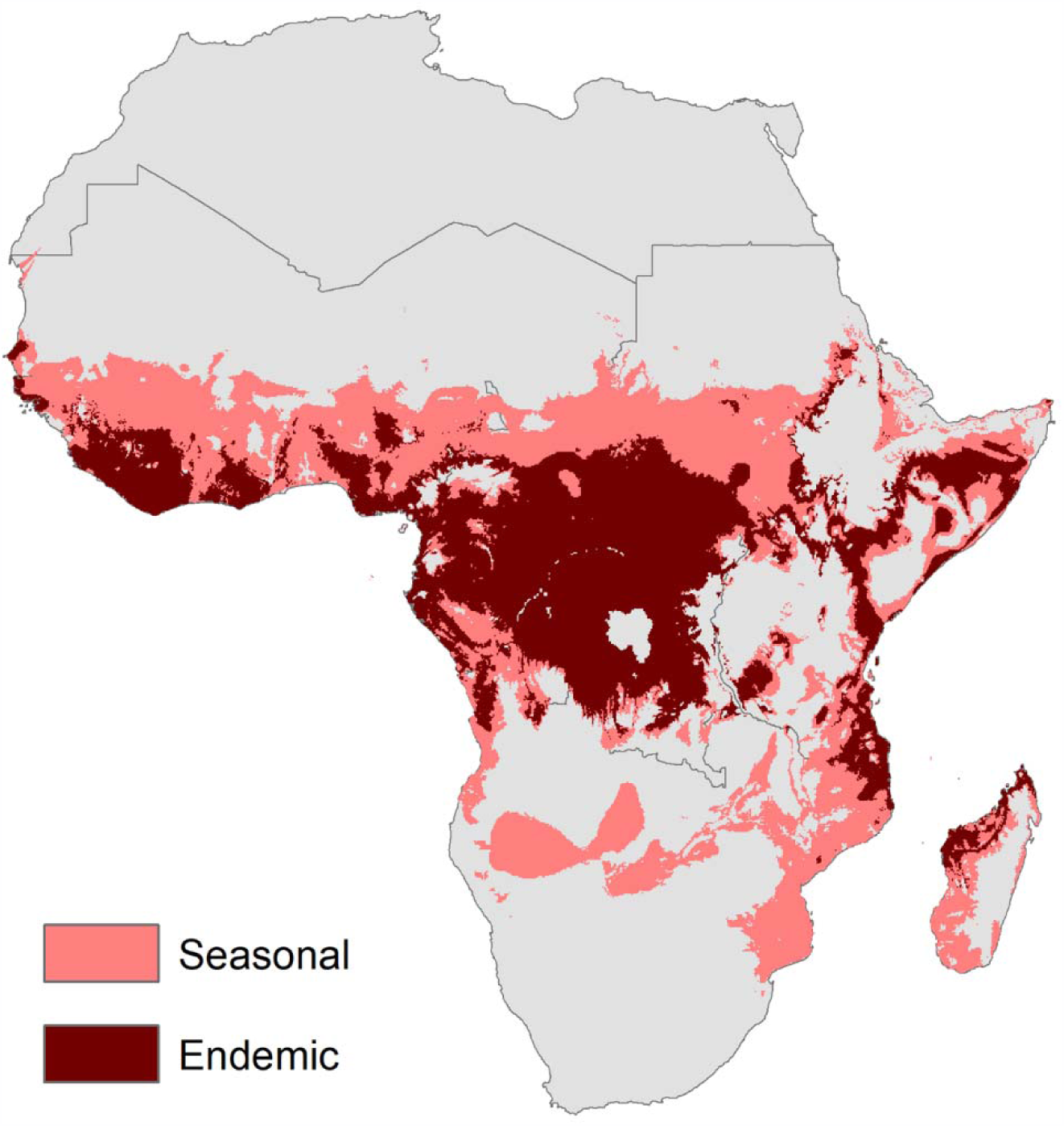
Modeled endemic (10-12 months) and seasonal (7-9 months) transmission suitability for malaria under current climate conditions.

The projected future climate impacts on malaria transmission suitability are shown for both RCP 4.5 and 8.5, for the three time horizons modeled, in Figure 3. Hotspots of endemic suitability will begin to emerge in the center of the continent, the East African highlands, the Lake Victoria region, and northern Zambia, becoming more pronounced in the latter part of the 21st century. A significant portion of these areas are located in Eastern Africa including Uganda, Kenya, and Tanzania, a region with currently lower suitability for endemic malaria transmission compared to Central and Western Africa. Additionally, areas predicted to have limited current suitability for *Anopheles* transmission may become seasonally suitable under conditions of a changing climate, including the Southern Africa region, which will see marked increases in areas suitable for seasonal and endemic malaria transmission (Figs. 2 and 3).

**Fig. 3.**
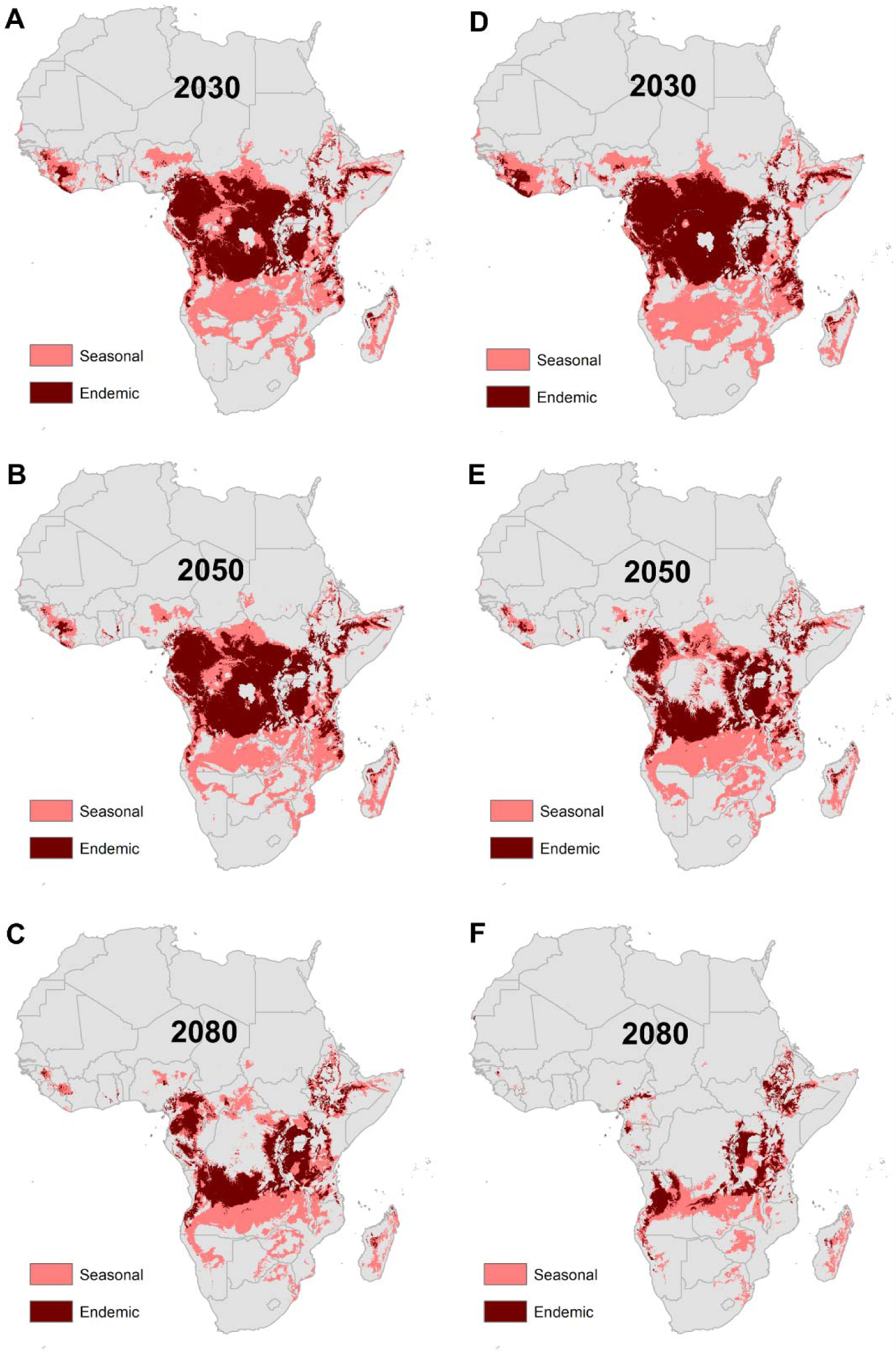
Modeled output of malaria transmission indicates shifting future endemic (dark red) and seasonal (light red) transmission suitability under two representative concentration pathways, RCP 4.5 (A, B, C) and RCP 8.5 (D, E, F), for the years 2030, 2050, and 2080.

Concentrated hotspots of seasonal suitability will begin to emerge in central Angola, northwestern Zambia, northern Tanzania, and the southern coast and northern part of Mozambique by 2030. This includes large portions of Zambia, Malawi, and Tanzania, eastern South Africa, Botswana, the highlands of Zimbabwe, northern Mozambique, and the Zambezi River Basin. Hotspots of seasonal malaria transmission suitability will either continue to concentrate, or will migrate both northward and southward into the highlands of Ethiopia and Southern Africa toward the latter part of the 21st century.

### Shifting burden of transmission suitability – people at risk

An additional 196–198 million people in Eastern and Southern Africa will be burdened with some degree of malaria transmission risk in the future due to shifting suitability by the 2080s. Regionally, by the year 2080 the worst-case scenario (RCP 8.5) places an additional 73.4 million people at risk from year-round exposure to transmission in Eastern Africa (Fig. 4). In spite of currently low endemic suitability, shifting seasonality in Southern Africa will place over 2.5 million additional people at risk for endemic transmission by the 2080s. In the short term, these changes are predicted to put the lives of 50.6–62.1 additional people at increased risk for endemic transmission, and 37.2–48.2 million people at risk for seasonal transmission, throughout Central, Eastern, and Southern Africa by the 2030s (Figs. 4 & 5). Given the strong empirical relationship between vector survival and temperature, as temperatures rise, exposure to malaria transmission is also expected to increase in previously unsuitable regions, such as those in the higher elevation regions of Southern and Eastern Africa. Countries likely to be impacted by these changes include northern Angola, southern DRC, western Tanzania, and central Uganda in 2030; by 2080 these changes will extend into western Angola, the upper Zambezi River Basin, and northeastern Zambia, and will become more concentrated along the East African highlands.

**Fig. 4.**
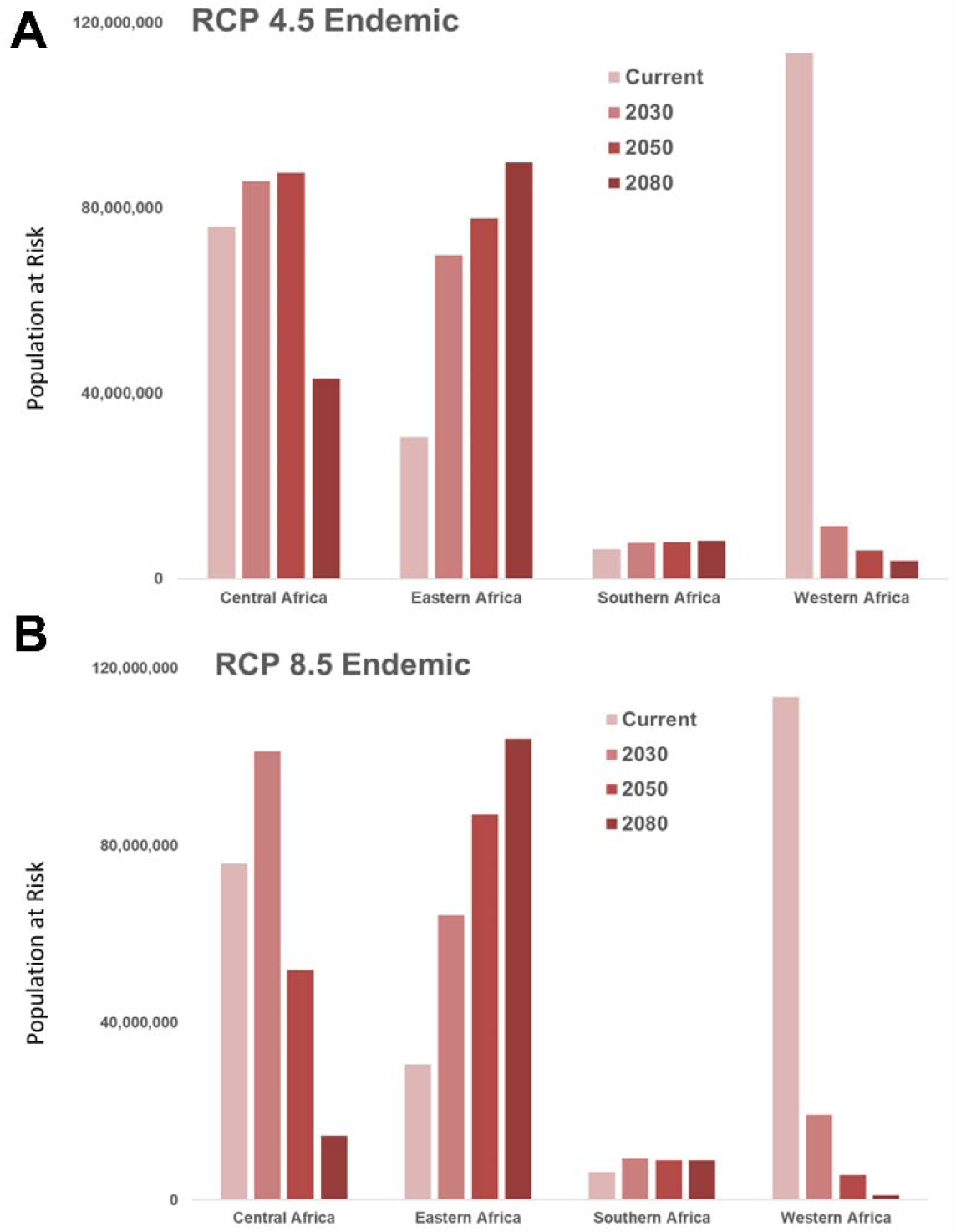
Population at risk (PAR) for exposure to endemic malaria transmission will change in the future as geographic suitability shifts under two scenarios of climate change, RCP 4.5 (A) and RCP 8.5 (B). Eastern Africa will regionally see dramatic increases PAR by the year 2080, while shifting suitability will largely relieve the burden of endemic transmission in Western Africa.

**Fig. 5.**
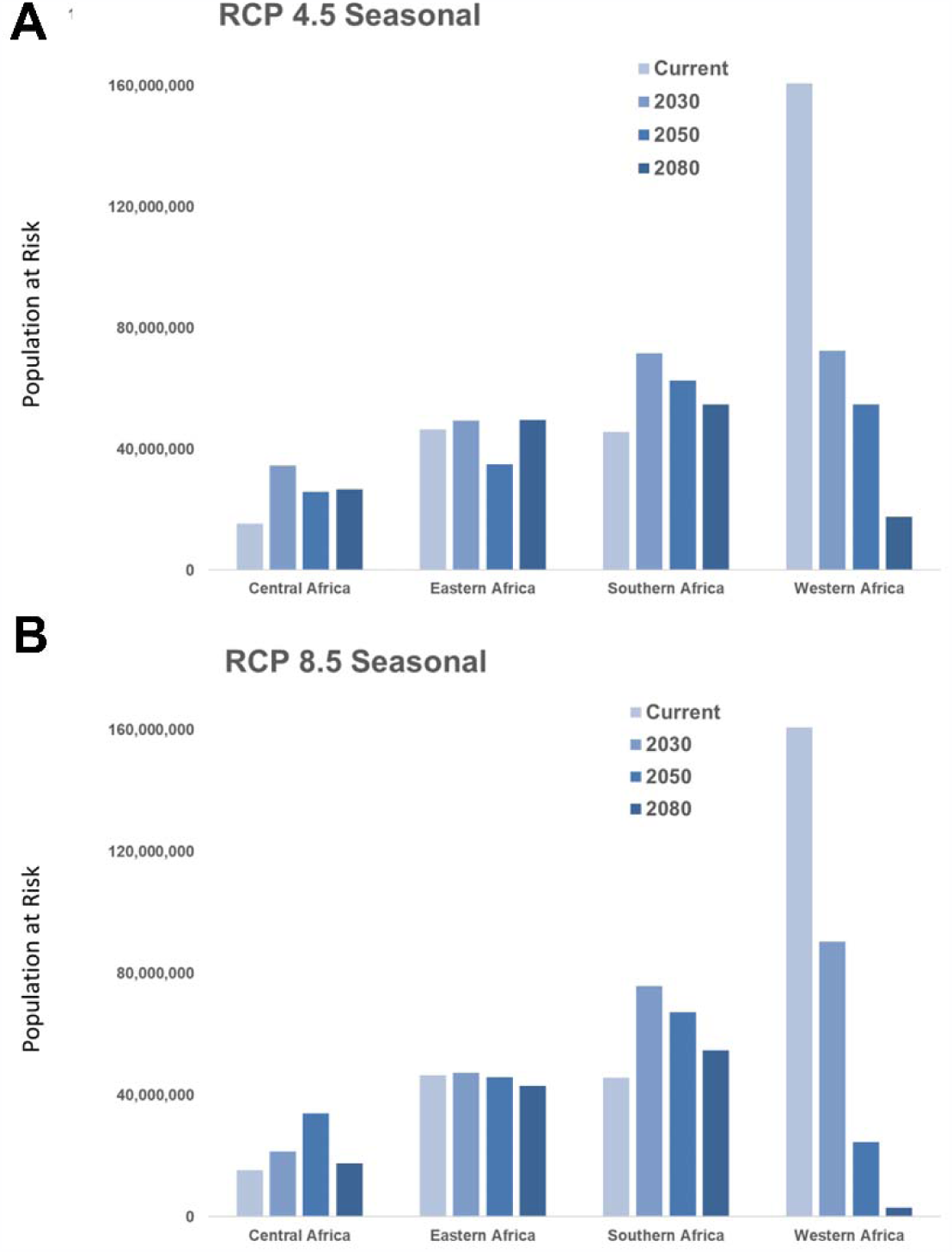
Population at risk (PAR) for exposure to seasonal malaria transmission will change in the future as geographic suitability shifts under two scenarios of climate change, RCP 4.5 (A) and RCP 8.5 (B). Southern Africa is predicted to have increased seasonal transmission, while shifting suitability will largely decrease seasonal transmission in Western Africa.

These shifts in the geographic range of malaria suitability, broadly consistent across both scenarios of future climate, suggest both decreases and increases in the number of people exposed, depending on the climate scenario. The geographic and temporal evolution of future suitability of areas for malaria-transmitting *Anopheles* mosquitoes is closely tied to expected temperature changes under both RCP scenarios (Fig. 3). As temperatures rise, even within the next 12 years (by 2030), important changes are anticipated. Shifting suitability due to climate change will place additional people at risk despite reductions endemic and seasonal malaria transmission, resulting in a net gain of 58.7 to 60.4 million people who experience some level of malaria risk in Western Africa by the 2030s. Large areas of coastal Western Africa and the Horn of Africa will likely exceed mosquitoes’ thermal tolerance, with suitability disappearing. At the same time, rising temperatures will likely increase the southern range of seasonal suitability for *Anopheles* mosquitoes into Southern and Central Africa, into western Tanzania. As temperatures continue to rise (2050s), both endemic and seasonal zones will continue to exhibit an eastward shift, with thermal threshold exceedance again apparent under the worst-case scenario (RCP 8.5), eliminating suitability across Central Africa. The end-of-the-century scenarios (2080) concentrate areas of endemism in previously unsuitable or marginally suitable areas, namely the highlands of East Africa and Southern Africa. Where the number of months of suitability for *Anopheles* survival decrease, opportunities will emerge to alter and define more targeted seasonal responses, either reducing the cost of interventions or providing a window into potential eradication to malaria exposure. Targets of opportunity include Central Africa (the Central African Republic, western Congo, Cameroon, and Equatorial Guinea) and coastal East Africa (Tanzania and Kenya).

### Novel Endemic and Seasonal Risk

Some parts of Sub-Saharan Africa currently predicted to experience no malaria transmission suitability risk will experience shifting suitability, resulting in novel areas with no history of malaria transmission becoming suitable for endemic and seasonal transmission in the future. As seen in Figure 6, for RCP 4.5, this exposes populations along an arc extending into East Africa, leading to dramatic PAR increases for regional exposures, particularly novel endemic exposure increase in East Africa, and novel seasonal exposures in Southern Africa (Figure 7).

**Figure 6:**
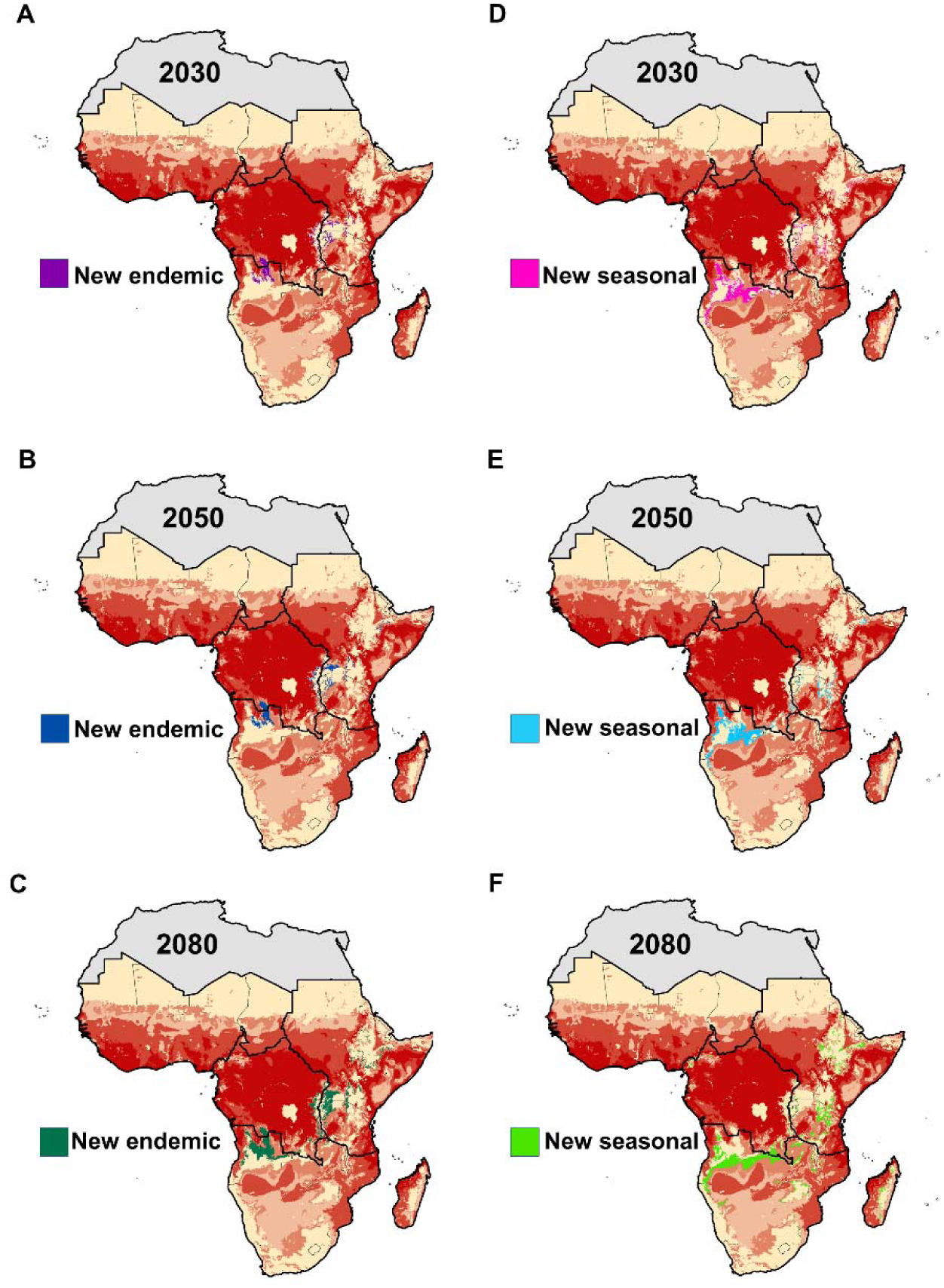
New areas of endemic (A-C) and seasonal (D-F) suitability, under RCP 4.5 for 2030, 2050, and 2080. Red shading intensity indicates current malaria suitability season.

**Figure 7:**
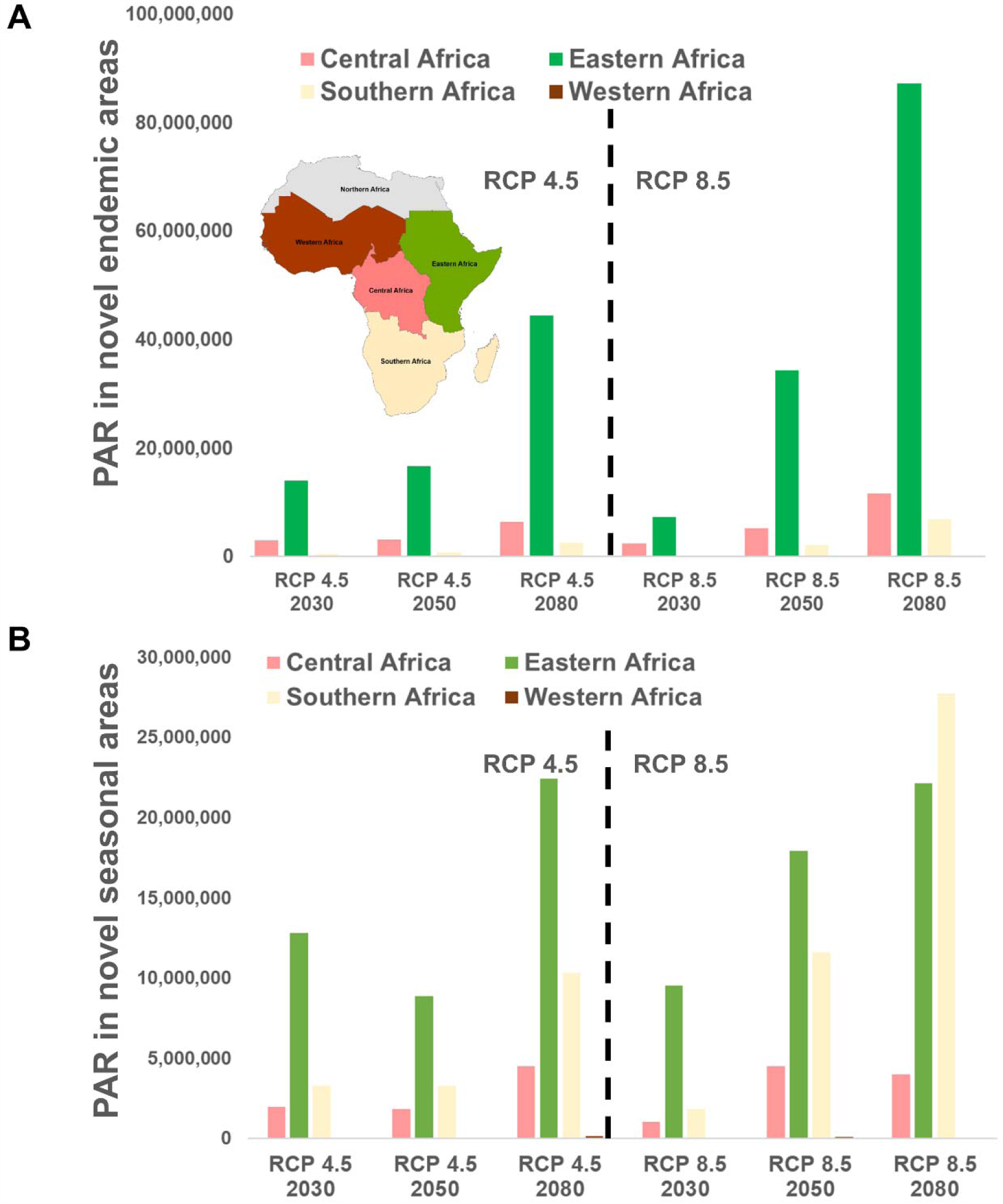
The number of people at risk (PAR) in A. newly endemic (10-12 month) suitable areas, and B. newly seasonal (7-9 month) suitable areas, for RCP 4.5 and RCP 8.5, in 2030, 2050, 2080

## Discussion

The changes in the geographic range of malaria suitability, broadly consistent across both scenarios of future climate, suggest that the number of people exposed to conditions of malaria suitability will both increase and decrease in Sub-Saharan Africa, depending on the region. Thus, as some populations experience reduced burden of malaria risk in the future, shifting suitability will increasingly place naïve populations at risk for outbreaks, particularly in Southern and Central Africa. Malaria outbreaks that occur where people have little or no immunity to the disease can lead to epidemic conditions, especially among vulnerable groups such as women and children [1,20]. This research identifies “hotspots” where current exposure, and therefore immunity, is nonexistent; these areas could see epidemic “flares” as climate conditions affect vector survival and reproduction. This effect may be further exacerbated in novel areas with no previous history of malaria exposure, where both immunity and knowledge regarding malaria prevention are lacking [21–23]. Malaria outbreaks occurring where people have acquired immunity due to prolonged and repeated malaria exposure trigger management actions employing a cadre of tools, including vector control and case management approaches to prevent or reduce transmission [23,24].

These results enable us to pinpoint regions where interventions need to be revisited to consider how climate will alter risk profiles in the future. The strong seasonal cycle of malaria across Southern Africa is related to climate and weather conditions [25,26]. Thus, during some periods of the year, climate conditions are not conducive to spread of the disease. Given the strong empirical relationship between vector survival and temperature, as temperatures rise exposure to malaria transmission is expected to increase in previously unsuitable regions, such as those in the higher elevation regions of Southern and East Africa. A key concern with climate change impacts is whether climate change will lengthen the period of the year during which diseases can establish and be transmitted. For example, areas where spring and autumn are now too cold for the reproduction of malaria vectors may become more suitable in the future. In these areas, increases in temperature may not impact midsummer malaria incidence greatly, but may result in a longer season, extending into both spring and autumn, during which malaria incidences will occur. In some cases, malaria may shift from being a seasonal disease burden to a year-round burden. This will necessitate different types of management and control interventions than those currently in place for short-season malaria [27,28]. Where the number of months of suitability for *Anopheles* survival decreases, opportunities will emerge to alter and define more targeted seasonal responses – either reducing the cost of interventions or providing a window into potential eradication to malaria exposure. An increase in the number of months where conditions are suitable for mosquito survival will require responses to be extended for longer periods of time, increasing resource needs (e.g. staff time, medicines) as well as costs [29]. In examining areas where malaria suitability is currently considered seasonally restricted, but will likely become more prevalent throughout the year, public health planners can anticipate which regions may require an extended investment pipeline.

A fundamental underpinning of modeling the response of vector-borne diseases to climate and ecology is the choice of model process. Previous approaches, such as that of the Malaria Atlas Project (MAP) and the Mapping Malaria Risk in Africa (MARA) project, are essentially top-down, wherein empirical data collected on the ground are matched to local climate conditions, and suitability established via geostatistical methods. In contrast, the modeling approach used here is mechanistic and “bottom-up,” wherein the life history of mosquitoes and pathogens, and their responses to temperature, are explicitly quantified based on empirical, laboratory-based data and incorporated into the model to predict where suitability for transmission is likely to occur. A mechanistic model, built independently of case outcome data, allows for validation with empirical, field-collected data, and obviates the bias of modeling data while intervention is ongoing, as is inevitably the case with previous approaches [30].

While substantial progress has been made in recent years in the provision and use of climate projections, considerable uncertainties remain with their use [31]. Using climate science research results to inform the decision process about which policies or specific measures are needed to tackle climate impacts requires acknowledging the uncertainties inherent in climate projections. These uncertainties may arise from mathematical reductions (parameterizations) of climate phenomena; potential socioeconomic technological pathways and attendant carbon cycle feedbacks that influence atmospheric concentrations of key greenhouse gases; imperfect scientific knowledge and the computational constraints of modelling regional detail while still incorporating relevant large-scale climate patterns; and the relationship between climate models and their relative impacts on key sectors and resources [31–33]. Furthermore, uncertainty can arise over the chance of a single event (for example, crossing a threshold), recurrent events (the return period of a flood, for example), discrete events (hurricane frequency), and complex events (for example, the interplay of different factors that lead to drought) [34]. Recognizing this, good practice is followed by incorporating a multimodel range of climate projections rather than a single model, as performed in this study [31,35,36]. For the population data specifically, it is important to recognize that the projected population for 2020 is used to calculate the numbers of people potentially affected by changing suitability conditions across all future time periods. As with climate models, these projections do not necessarily capture all of the factors that drive population movement and growth and should be taken as best modelled estimates rather than exact values.

The study results are based on the temperature response curves of both *Anopheles* mosquitoes and malaria pathogens. Nevertheless, many studies point to the critical role that rainfall plays in vector survival across Sub-Saharan Africa [12,14,15]. For example, single, intense rainfall events can wash away critical breeding sites, leading to a reduction in transmission potential [16,37]. Similarly, too little rainfall can limit mosquito survival as moisture is a prerequisite for breeding habitat [38]. The approach herein addresses this second issue by masking out areas that are too arid for mosquito survival. While the relationship between rainfall and *Anopheles* survival is critical, the available projections of rainfall are uncertain at the geographic scale of this work and therefore are not considered in this analysis.

Geographically projected model outputs are a useful component of a planning and intervention framework, providing a means of communicating key areas of risk and affected populations to decisionmakers. Anticipation of not only the location and time, but the duration of potential outbreak events will facilitate the development of efficient and timely agency responses. Moreover, this framework serves as a foundation for scenario analysis, explicitly modeling risk of exposure for different climate scenarios and time horizons. The range of potential outcomes allows governments and agencies the flexibility needed to reasonably anticipate resource use and funding needs, enabling the development of adaptive intervention strategies for both near and long-term outcomes.

## Conclusions

Addressing the changing risk profiles projected in this suitability analysis will require modifying current interventions and programs and implementing new ones to explicitly consider climate variability and change. Opportunities for improved responses also exist, including detailed geographic targeting, optimizing strategies and seasonal alignment with interventions. Identifying high risks in new areas of suitability present opportunities for informed action. Where malaria suitability is currently nonexistent to newly suitable, whether seasonal or endemic, the risks are critical, especially given that local populations’ immunity will be low. This could lead to the potential emergence of novel strains, rapid resistance, and untimely identification, translating into epidemic outbreaks. To respond, targeted and informed geographic surveillance in these regions could help to prepare timely responses before epidemic outbreaks occur. Knowing where and when more people will potentially be exposed offers an opportunity to increase the investment timeframe (seasonal to year-round), optimize vector control, and improve case management, with the evidence base to support these actions. Moving down the path toward elimination for some regions, where malaria transmission suitability decreases, opportunities will arise to focus resources on making surveillance and response systems increasingly sensitive and focused to identify, track, and respond to malaria cases and any remaining transmission foci.

## Abbreviations

MAP: Malaria Atlas Project
MARA: Mapping Malaria Risk in Africa
NDVI: normalized difference vegetation index
GCM: global climate model
CMIP5: Coupled Model Intercomparison Project
CF: change factor
RCP: representative concentration pathway
PAR: population at risk
GPW: Gridded Population of the World

## Declarations

## Availability of Data and Materials

Data sharing is not applicable to this article as no datasets were generated or analysed during the current study.

## Competing Interests

The authors declare no competing interests

## Consent for Publication

Not applicable

## Ethics Approval and Consent to Participate

Not applicable

## Funding

This analysis was funded by by the United States Agency for International Development through the Adaptation Thought Leadership and Assessments (ATLAS) Task Order No. AID-OAA-I-14-00013, under the Restoring the Environment through Prosperity, Livelihoods, and Conserving Ecosystems (REPLACE) IDIQ.

## Authors’ Contributions

SJR and FZ conceived of the study, SJR ran analyses, FZ, SJR, and CAL wrote, edited, and refined the manuscript

## Acknowledgements

The authors would like to thank Tegan Blaine and Colin Quinn of USAID’s Africa bureau for their guidance in aligning the assessment to on the ground management decisions; and Jordan Burns and Rene Salgado of the President’s Malaria Initiative for the review and comments.

## References

1. World Health Organization. World Malaria Report 2018. 2018 Nov p. 210.

2. Ryan SJ, McNally A, Johnson LR, Mordecai EA, Ben-Horin T, Paaijmans K, et al. Mapping Physiological Suitability Limits for Malaria in Africa Under Climate Change. Vector-Borne and Zoonotic Diseases. 2015;15:718–25.

3. Mordecai EA, Paaijmans KP, Johnson LR, Balzer C, Ben-Horin T, de Moor E, et al. Optimal temperature for malaria transmission is dramatically lower than previously predicted. Thrall P, editor. Ecology Letters. 2013;16:22–30.

4. Pascual M, Ahumada JA, Chaves LF, Rodo X, Bouma M. Malaria resurgence in the East African highlands: temperature trends revisited. National Acad Sciences; 2006.

5. Siraj AS, Santos-Vega M, Bouma MJ, Yadeta D, Carrascal DR, Pascual M. Altitudinal Changes in Malaria Incidence in Highlands of Ethiopia and Colombia. Science. 2014;343:1154–8.

6. Tanser FC, Sharp B, le Sueur D. Potential effect of climate change on malaria transmission in Africa. The Lancet. 2003;362:1792–8.

7. Gething PW, Smith DL, Patil AP, Tatem AJ, Snow RW, Hay SI. Climate change and the global malaria recession. Nature. 2010;465:342–5.

8. Gething P, Van Boeckel T, Smith D, Guerra C, Patil A, Snow R, et al. Modelling the global constraints of temperature on transmission of Plasmodium falciparum and P. vivax. Parasites & Vectors. 2011;4:92.

9. Binka F, De Savigny D. MONITORING FUTURE IMPACT ON MALARIA BURDEN IN SUB-SAHARAN AFRICA. The American Journal of Tropical Medicine and Hygiene. 2004;71:224–31.

10. Hijmans RJ, Cameron SE, Parra JL, Jones PG, Jarvis A. Very high resolution interpolated climate surfaces for global land areas. International Journal of Climatology. 2005;25:1965–78.

11. Navarro Racines CE, Tarapues Montenegro JE, Thornton P, Jarvis A, Ramirez Villegas J. CCAFS-CMIP5 Delta Method Downscaling for monthly averages and bioclimatic indices of four RCPs [Internet]. World Data Center for Climate (WDCC) at DKRZ; 2019 [cited 2019 May 2]. Available from: http://cera-www.dkrz.de/WDCC/ui/Compact.jsp?acronym=CCAFS-CMIP5_downscaling

12. Thomson MC, Mason SJ, Phindela T, Connor SJ. Use of rainfall and sea surface temperature monitoring for malaria early warning in Botswana. Am J Trop Med Hyg. 2005;73:214–21.

13. Grover-Kopec E, Kawano M, Klaver RW, Blumenthal B, Ceccato P, Connor SJ. An online operational rainfall-monitoring resource for epidemic malaria early warning systems in Africa. Malar J. 2005;4:6.

14. Pascual M, Cazelles B, Bouma MJ, Chaves LF, Koelle K. Shifting patterns: malaria dynamics and rainfall variability in an African highland. Proc Biol Sci. 2008;275:123–32.

15. Craig MH, Snow RW, le Sueur D. A climate-based distribution model of malaria transmission in sub-Saharan Africa. Parasitol Today (Regul Ed). 1999;15:105–11.

16. Paaijmans KP, Wandago MO, Githeko AK, Takken W. Unexpected High Losses of Anopheles gambiae Larvae Due to Rainfall. Carter D, editor. PLoS ONE. 2007;2:e1146.

17. Suzuki R, Xu J, Motoya K. Global analyses of satellite-derived vegetation index related to climatological wetness and warmth. International Journal of Climatology. 2006;26:425–38.

18. US Geological Survey and US Agency for International Development. FEWS-NET (Famine Early Warning Systems Network) [Internet]. 2018 [cited 2018 Jan 19]. Available from: https://earlywarning.usgs.gov/fews/search/Africa

19. Center For International Earth Science Information Network-CIESIN-Columbia University. Gridded Population of the World, Version 4 (GPWv4): Population Density Adjusted to Match 2015 Revision of UN WPP Country Totals [Internet]. Palisades, NY: NASA Socioeconomic Data and Applications Center (SEDAC); 2016 [cited 2018 Mar 8]. Available from: http://beta.sedac.ciesin.columbia.edu/data/set/gpw-v4-population-density-adjusted-to-2015-unwpp-country-totals

20. Trape J-F, Rogier C. Combating malaria morbidity and mortality by reducing transmission. Parasitology Today. 1996;12:236–40.

21. Ndyomugyenyi R, Magnussen P, Clarke S. Malaria treatment-seeking behaviour and drug prescription practices in an area of low transmission in Uganda: implications for prevention and control. Transactions of the Royal Society of Tropical Medicine and Hygiene. 2007;101:209–15.

22. Doolan DL, Dobano C, Baird JK. Acquired Immunity to Malaria. Clinical Microbiology Reviews. 2009;22:13–36.

23. Kiszewski AE, Teklehaimanot A. A review of the clinical and epidemiologic burdens of epidemic malaria. Am J Trop Med Hyg. 2004;71:128–35.

24. Abeku TA. Response to Malaria Epidemics in Africa. Emerging Infectious Diseases. 2007;13:681–6.

25. Adeola A, Botai J, Rautenbach H, Adisa O, Ncongwane K, Botai C, et al. Climatic Variables and Malaria Morbidity in Mutale Local Municipality, South Africa: A 19-Year Data Analysis. International Journal of Environmental Research and Public Health. 2017;14:1360.

26. Ikeda T, Behera SK, Morioka Y, Minakawa N, Hashizume M, Tsuzuki A, et al. Seasonally lagged effects of climatic factors on malaria incidence in South Africa. Scientific Reports [Internet]. 2017 [cited 2019 Sep 20];7. Available from: http://www.nature.com/articles/s41598-017-02680-6

27. Walker PGT, Griffin JT, Ferguson NM, Ghani AC. Estimating the most efficient allocation of interventions to achieve reductions in Plasmodium falciparum malaria burden and transmission in Africa: a modelling study. The Lancet Global Health. 2016;4:e474–84.

28. World Health Organization. WHO Policy Recommendation: Seasonal Malaria Chemoprevention (SMC) for Plasmodium falciparum malaria control in highly seasonal transmission areas of the Sahel sub-region in Africa [Internet]. Global Malaria Program, World Health Organization; 2012. Available from: https://www.who.int/malaria/publications/atoz/smc_policy_recommendation_en_032012.pdf?ua=1

29. Goodman C, Coleman P, Mills A. Cost-effectiveness of malaria control in sub-Saharan Africa. The Lancet. 1999;354:378–85.

30. Mordecai EA, Caldwell JM, Grossman MK, Lippi CA, Johnson LR, Neira M, et al. Thermal biology of mosquitoLborne disease. Byers J (Jeb), editor. Ecology Letters. 2019;22:1690–708.

31. Knutti R, Sedláček J. Robustness and uncertainties in the new CMIP5 climate model projections. Nature Climate Change. 2013;3:369–73.

32. Raäisaänen J. How reliable are climate models? Tellus A: Dynamic Meteorology and Oceanography. 2007;59:2–29.

33. Knutti R, Allen MR, Friedlingstein P, Gregory JM, Hegerl GC, Meehl GA, et al. A Review of Uncertainties in Global Temperature Projections over the Twenty-First Century. Journal of Climate. 2008;21:2651–63.

34. Palmer TN, Shutts GJ, Hagedorn R, Doblas-Reyes FJ, Jung T, Leutbecher M. REPRESENTING MODEL UNCERTAINTY IN WEATHER AND CLIMATE PREDICTION. Annual Review of Earth and Planetary Sciences. 2005;33:163–93.

35. Knutti R, Abramowitz G, Collins M, Eyring V, Glecker P, Hewitson B, et al. Good Practice Guidance Paper on Assessing and Combining Multi Model Climate Projections. IPCC Working Group I Technical Support Unit; 2010.

36. Meehl GA, Covey C, Delworth T, Latif M, McAvaney B, Mitchell JFB, et al. THE WCRP CMIP3 Multimodel Dataset: A New Era in Climate Change Research. Bulletin of the American Meteorological Society. 2007;88:1383–94.

37. Zermoglio F, Ryan SJ, Swaim M. Shifting burdens: malaria risk in a hotter Africa. USAID; 2019.

38. Charlwood JD, Kihonda J, Sama S, Billingsley PF, Hadji H, Verhave JP, et al. The rise and fall of *Anopheles arabiensis* (Diptera: Culicidae) in a Tanzanian village. Bulletin of Entomological Research. 1995;85:37–44.

